# Pivotal role of biallelic frequency analysis in identifying copy number alterations using genome-wide methods in tumors with a high level of aneuploidy

**DOI:** 10.1101/2024.03.14.584989

**Authors:** Julia Rymuza, Renata Woroniecka, Beata Grygalewicz, Mateusz Bujko

## Abstract

Chromosome number abnormalities is one of the hallmarks of cancer. DNA copy number alterations (CNA) are studied using various genome-wide methods. In our study we investigated CNA in human pituitary tumors using three platforms CytoSNP-850K microarrays, low-pass whole-genome sequencing (average x7 coverage, LPWGS), and Infinium Methylation EPIC array. Virtual karyotypes based on each dataset were generated using open-source software packages for each sample. Concordant CNA profiles were found for most of tumor. Surprisingly, substantial discrepancies between results from SNP arrays and LPWGS/EPIC arrays were identified in 20% of tumors, for which discrimination of true karyotype was required. B-allelic frequency data from SNP arrays was crucial to adjust normal ploidy level as ultimately verified with FISH. The discrepancy between virtual karyotypes was more pronounced the more CNAs were found. When CNAs covered more than half of genome the level of normal/diploid copy number was incorrectly set with methods, based solely on signal intensity/read-counts coverage. To conclude, CNA analysis with methods such as LPWGS and methylation arrays in highly aneuploid tumors are prone to a bias from improper normal ploidy level setting. These methods are commonly used therefore we aimed to aware the scientific community about this underestimated methodological problem.

---

Aneuploidy is a copy number alteration (CNA) that encompasses a wide spectrum of changes ranging from single arm gains/losses to complex rearrangements involving multiple chromosomes up to whole genome duplication [1]. Although the role of aneuploidy in carcinogenesis has been a subject of research since its discovery in the 19th century, its mechanisms remain unclear [2, 3]. It is estimated to occur in 90% of human cancer, with different prevalence between tumor types [4–6].

Nowadays, various laboratory methods for studding whole genome CNA in cancer are available including: SNP microarray, whole genome sequencing (WGS) (including low-coverage WGS), and DNA methylation microarrays [7]. These methods are cost-effective and widely used for both scientific and clinical applications. Although they can be considered equivalent, they are based on distinct technical and analytical principles. Analysis based on SNP array includes two types of signals: B-allele frequency (BAF) and signal intensity logR ratio. There are different approaches to analyzing these data, with some optimized to work with cancer samples [8, 9]. Inferring CNA from WGS data can be done based on the profile of read depth (RD) of aligned reads [10]. The difference in coverage of chromosomal regions by sequencing reads indicates copy number alteration. The advantage of WGS is that low sequencing coverage was recognized to be efficient for copy number detection [10]. In turn, DNA methylation microarrays are commonly used for profiling genome-wide CNA because they allow for both epigenetic and cytogenetic profiling of the sample in one procedure. The most common tool for analysis of CNA from DNA methylation microarray data is conumme.

Recent work from our group focuses on molecular profiling on somatotroph pituitary neuroendocrine tumors (sPitNET), which are nonmalignant intercranial tumors originating from the cells of anterior pituitary. In our study, we used three distinct methods that allow inferring whole-genome copy number profile: Illumina’s beadChip technology CytoSNP-850K genotyping oligonucleotide microarrays (Illumina), low pass WGS (pair-end 2 × 150 bp Illumina sequencing with an average x7 coverage) and Infinium MethylationEPIC BeadChip Array (Illumina). These three platforms were used to generate virtual karyotypes using custom Python and R scripts for each tumor sample (as exampled in Fig. 1). In nearly one third of the samples (13/40), we observed surprising discrepancies between the results of the SNP array and those of the WGS and EPIC array (as exampled in Fig. 2). We found that discordance between results was related to genome instability load in the tumors and increased with the number of altered chromosomes. Since a standard, commonly used analytical approaches were used for CNA detection, with each method, we recognized the observed discrepancy as a symptom of the general problem. Our study included samples with an apparently large spectrum of aneuploidy (ranging from 0 to 91% of the genome affected by CNA); therefore, we made an attempt to reveal the source of these discrepancies by exploring how different levels of aneuploidy influence the computational analysis.

**Figure 1.**
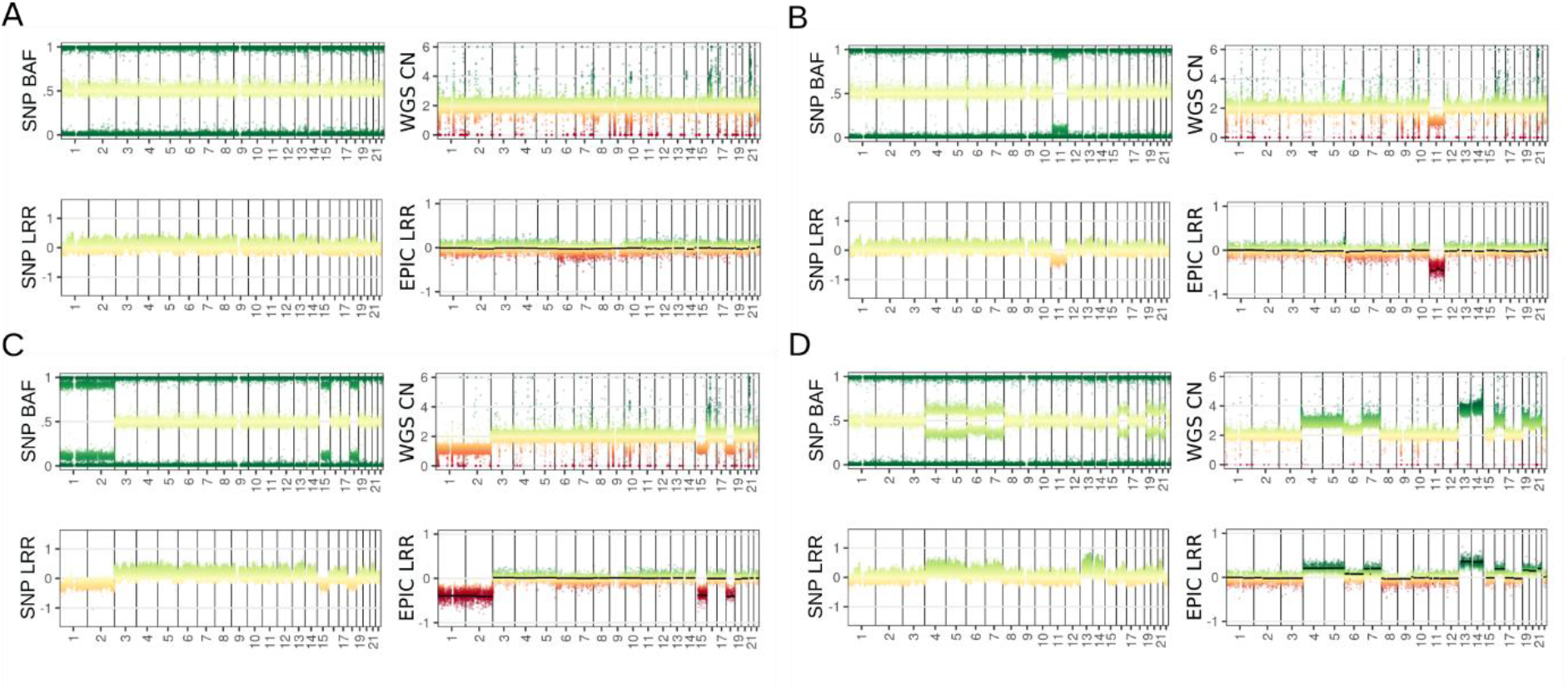
Virtual karyotypes of samples with low level of aneuploidy. The baseline copy number of the sample for the WGS data is two, and for the EPIC array it is 0. A-D) Stable samples, with concordant results from SNP array, WGS, and EPIC array. SNP BAF - B-allele frequency from SNP array; SNP LRR - signal intensity logR ratio from SNP array; WGS CN – normalized read depth calculated by CNVpytor from WGS; EPIC LRR - ratio of normalized intensities calculated by conumee from EPIC array.

**Figure 2.**
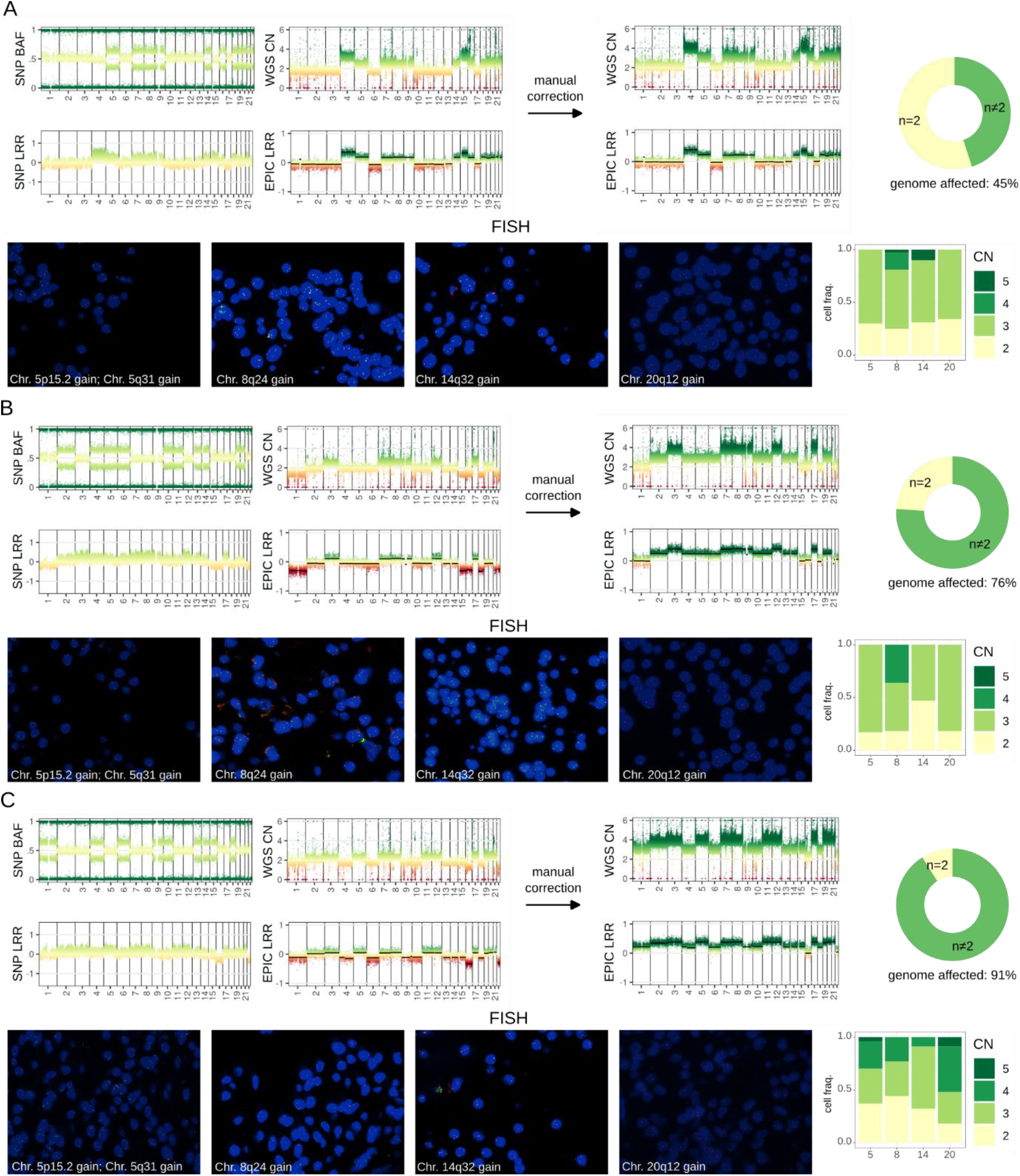
Virtual karyotypes of samples with high levels of aneuploidy. The baseline copy number of the sample for the WGS data is two, and for the EPIC array it is 0. A-B) Highly aneuploid samples with diverging results from WGS and EPIC array from results from SNP array. With manually corrected signal intensities for WGS and EPIC array to match baseline to two copies of chromosomes and information about the fraction of genome affected by CNA. And results from FISH representing cell fractions having given number of copies of analyzed chromosomes. SNP BAF - B-allele frequency from SNP array; SNP LRR - signal intensity logR ratio from SNP array; WGS CN – normalized read depth calculated by CNVpytor from WGS; EPIC LRR- ratio of normalized intensities calculated by conumee from EPIC array.

Virtual karyotypes for each sample were created by plotting following data: B-allele frequency (BAF) and signal intensity logR ratio (LRR) from SNP array data processed with GenomeStudio (Illumina); read depth (RD) from WGS data calculated using CNVpytor [11]; and ratio of normalized intensities (LRR EPIC) from EPIC array, which was acquired using R conumee [12], with segments from the results from MolecularNeuropathology.org [13]. Signal intensity measures: LRR from SNP and EPIC arrays and RD from WGS directly show changes in chromosome copy number within the sample while patterns of BAF from SNP array distinguish between chromosomes with even and odd numbers of copies.

We analyzed samples with low level of aneuploidy including stable genome (Fig. 1A), single chromosome/chromosomal arm deletions (Fig. 1B) as well as deletion (Fig. 1C) and duplication (Fig. 1D) of multiple chromosomes. Full concordance between results of all three data types (between virtual karyotypes), including LRR from the SNP array, RD from WGS, and LRR from EPIC, was observed for the samples with low level of aneuploidy. Deletions appeared as lower signal level (Fig. 1B, C) while duplications had a higher signal level (Fig. 1D).

However, with the increasing level of aneuploidy in a sample (affecting more than half of the genome), we observe a discrepancy between BAF signal and signal intensity methods (RD from WGS and LRR from EPIC array). For the sample with intermediate aneuploidy (duplication of 13 chromosomes, Fig. 2A) the baseline for signal intensity methods no longer corresponds to a specific chromosome copy number confusing the interpretation of the results. In sample with high aneuploidy level (duplication of 18 chromosomes, Fig. 2B), chromosomes with baseline level from signal intensity methods have inconsistently an odd number of copies according to BAF from SNP array. Similarly, in a sample with very high aneuploidy (duplication of 20 chromosomes, Fig. 2C) chromosomes signal level slightly below baseline from signal intensity methods also have an odd number of copies based on BAF from SNP array. The chromosome with the lowest copy number based on signal intensity methods has an even number of copies according to BAF track. For both this samples and for the other 6 included in our study, chromosomes with normal ploidy according to SNP arrays are recognized by algorithms based on WGS and methylation data as monosomes, while those chromosomes with an additional copy (n=3) are recognized as diploids (n = 2).

Information on the SNP distribution from SNP microarray appears to be pivotal in determining which is the true karyotype, since it unequivocally shows even or odd number of chromosomes. Thus, BAF profile do not discriminate between two and four copies of a chromosome but clearly distinguish 2 and 3 copies as well as 2 and 1 copies.

The WGS and EPIC-based karyotypes can be adjusted to match the information represented by the BAF (Fig. 2). This adjustment, which is actually a manual shift of the calculated values in WGS and EPIC-based karyotypes, results in complete overlap between copy number profiles obtained with three independent methods. The real CNA profile was verified in our study by FISH in FFPE tissue using the probes for selected loci on chromosomes 5p15.2, 5q31, 8q24, 14q32, 20q12 that were commonly duplicated in a group of tumors with a high level of ploidy. Microscopic evaluation with quantification by signal counts definitely confirmed the validity of the adjustment of WGS and EPIC-based karyotypes (Fig. 2).

Our observations highlight the challenges in analyzing samples with unknown ploidy. Baseline of normalized signal intensity usually represents the average ploidy of the sample. When the analyzed sample is mostly diploid, this approach allows accurate identification of chromosomal deletions and duplication. However, with increasing aneuploidy level, the calculated baseline is more and more biased, and may lead to an incorrect interpretation of the data. In such a case, the baseline does not correspond to chromosomes with two copies and methods incorporating the BAF profile should be used. Importantly, it is not available for all the genome-wide methods including methylation microarrays.

Our result points pitfalls of using approaches that assumes that the examine sample is mainly diploid. It shows the need to visually inspecting data and the importance of including BAF signal to decode the number of copies. Beside whole genome approaches, the data from whole exome or genes panels sequencing are also commonly used for inferring the CNA pattern [7]. These methods allow for BAF analysis, and our experience highlights the need of including analysis of allelic distribution for reliable inferring CNA, if its available.

On the other hand, our results suggest a caution when interpreting that data from signal intensity/coverage depth only. In many already published studies, CNA was inferred from DNA methylation microarrays, at the margins of methylation profiling, where the baseline level was not fully controlled. For example: conumee package is incorporated into wildly used tool for the classification of central nervous system tumors based on methylation data [13]. This classifier is a fantastic and advanced tool for categorizing the samples that also provide a methylation array-based karyotype as a supplement to a final report. When our highly aneuploid samples were uploaded to this classifier, we obtained biased, mistaken karyotypes (Fig. 2) that required BAF-based adjustment. The potential sensitivity of this tool for normalization bias was already mentioned by the classifier authors [14] and our results are clear practical illustration of this problem.

The aim of this communication is to make the scientific community aware of this underestimated problem. In research the aneuploidy status of the sample is commonly unknown, and the possibility of potential normalization bias should be considered when planning a study and selecting a particular analytical method. Results of the previous studies (especially those on cancer genetics) where methods fragile to such a bias were used should also be treated with caution. Our data are in no way conclusive for estimating how the identified problem affects general knowledge in the field. However, working on pituitary tumors we noted a discrepancies between published results of CNA profiling and we assume that normalization problem could explain some of them.

## AUTHOR CONTRIBUTIONS

Conceptualization: JR, MB

Methodology: JR, MB, BG

Investigation: JR, MB, RW, BG

Visualization: JR, RW

Funding acquisition: MB

Project administration: MB

Supervision: MB

Writing – JR, MB

Writing – review & editing: RW, BG

## RESEARCH FUNDING

This work was supported by Maria Sklodowska-Curie National Research Institute of Oncology, grant SN/GW04/2020.

## REFERENCES

1. Lau TY, Poon RYC. Whole-Genome Duplication and Genome Instability in Cancer Cells: Double the Trouble. Int J Mol Sci 2023;24:3733.

2. Watkins TBK, Lim EL, Petkovic M, Elizalde S, Birkbak NJ, Wilson GA, et al. Pervasive chromosomal instability and karyotype order in tumour evolution. Nature;587. Epub ahead of print 2020. DOI:10.1038/s41586-020-2698-6.

3. Vasudevan A, Schukken KM, Sausville EL, Girish V, Adebambo OA, Sheltzer JM. Aneuploidy as a promoter and suppressor of malignant growth. Nature Reviews Cancer;21. Epub ahead of print 2021. DOI:10.1038/s41568-020-00321-1.

4. Knouse KA, Davoli T, Elledge SJ, Amon A. Aneuploidy in Cancer: Seq-ing Answers to Old Questions. Annu Rev Cancer Biol 2017;1:335–354.

5. Taylor AM, Shih J, Ha G, Gao GF, Zhang X, Berger AC, et al. Genomic and Functional Approaches to Understanding Cancer Aneuploidy. Cancer Cell;33. Epub ahead of print 2018. DOI:10.1016/j.ccell.2018.03.007.

6. Shih J, Sarmashghi S, Zhakula-Kostadinova N, Zhang S, Georgis Y, Hoyt SH, et al. Cancer aneuploidies are shaped primarily by effects on tumour fitness. Nature 2023;619:793–800.

7. Gordeeva V, Sharova E, Arapidi G. Progress in Methods for Copy Number Variation Profiling. International Journal of Molecular Sciences;23. Epub ahead of print 2022. DOI:10.3390/ijms23042143.

8. Popova T, Manié E, Stoppa-Lyonnet D, Rigaill G, Barillot E, Stern MH. Genome Alteration Print (GAP): A tool to visualize and mine complex cancer genomic profiles obtained by SNP arrays. Genome Biol;10. Epub ahead of print 2009. DOI:10.1186/gb-2009-10-11-r128.

9. Sun W, Wright FA, Tang Z, Nordgard SH, Van Loo P, Yu T, et al. Integrated study of copy number states and genotype calls using high-density SNP arrays. Nucleic Acids Res;37. Epub ahead of print 2009. DOI:10.1093/nar/gkp493.

10. Chaubey A, Shenoy S, Mathur A, Ma Z, Valencia CA, Reddy Nallamilli BR, et al. Low-Pass Genome Sequencing: Validation and Diagnostic Utility from 409 Clinical Cases of Low-Pass Genome Sequencing for the Detection of Copy Number Variants to Replace Constitutional Microarray. Journal of Molecular Diagnostics;22. Epub ahead of print 2020. DOI:10.1016/j.jmoldx.2020.03.008.

11. Suvakov M, Panda A, Diesh C, Holmes I, Abyzov A. CNVpytor: a tool for copy number variation detection and analysis from read depth and allele imbalance in whole-genome sequencing. Gigascience;10. Epub ahead of print 18 November 2021. DOI:10.1093/gigascience/giab074.

12. Hovestadt V, Zapatka M. conumee: Enhanced copy-number variation analysis using Illumina DNA methylation arrays. http://bioconductor.org/packages/conumee/.,. (2015, accessed 21 February 2024).

13. Capper D, Jones DTW, Sill M, Hovestadt V, Schrimpf D, Sturm D, et al. DNA methylation-based classification of central nervous system tumours. Nature 2018;555:469–474.

14. Capper D, Stichel D, Sahm F, Jones DTW, Schrimpf D, Sill M, et al. Practical implementation of DNA methylation and copy-number-based CNS tumor diagnostics: the Heidelberg experience. Acta Neuropathol;136. Epub ahead of print 2018. DOI:10.1007/s00401-018-1879-y.

